## Supplemental Files for "Regional Gene Expression in the Retina, Optic Nerve Head, and Optic Nerve of Mice with Experimental Glaucoma and Optic Nerve Crush"

Figures S1-8

Tables S1-3

*Spreadsheets S1-6 (as excel files)*

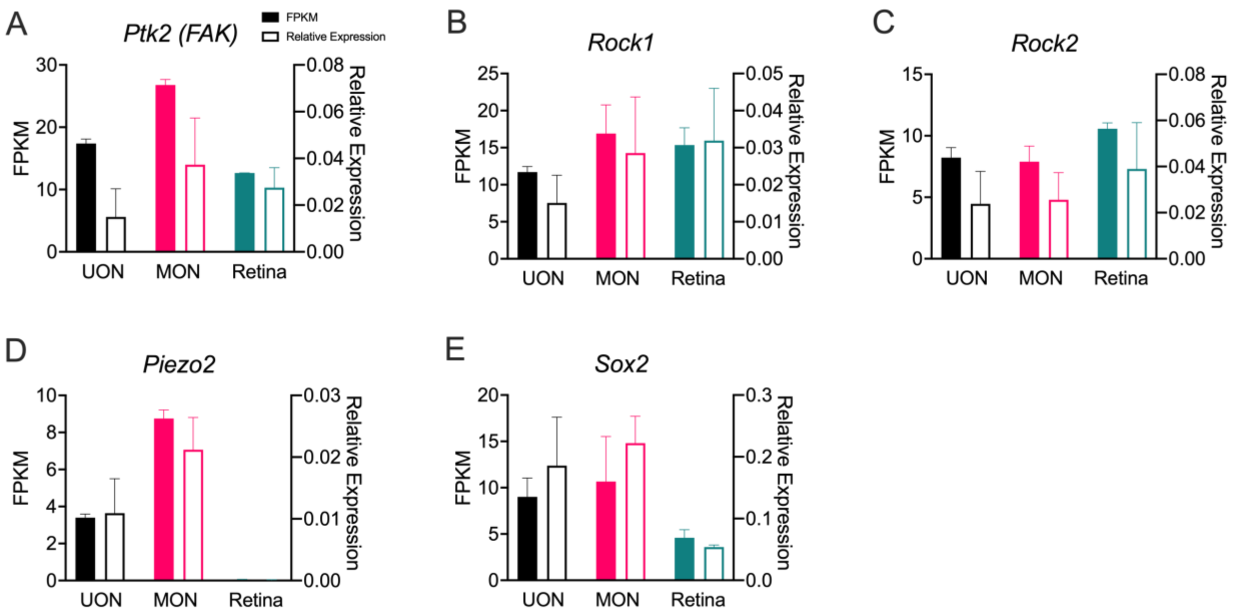


**Figure S1.** qPCR validation of naïve RNA-seq data. **(A-E)** Expression of *Ptk2* (A), *Rock1* (B), *Rock2* (C), *Piezo2* (D), and *Sox2* (E) in three naïve tissue regions: UON, MON, and retina. Left y-axis and filled bars represent FPKM (RNA-seq), while right y-axis and empty bars indicate relative expression via qPCR of independent tissue samples. Error bars indicate standard deviation. For RNA-seq, n = 2 (pooled) samples per tissue type. For qPCR, n = 6 individual samples per tissue group.

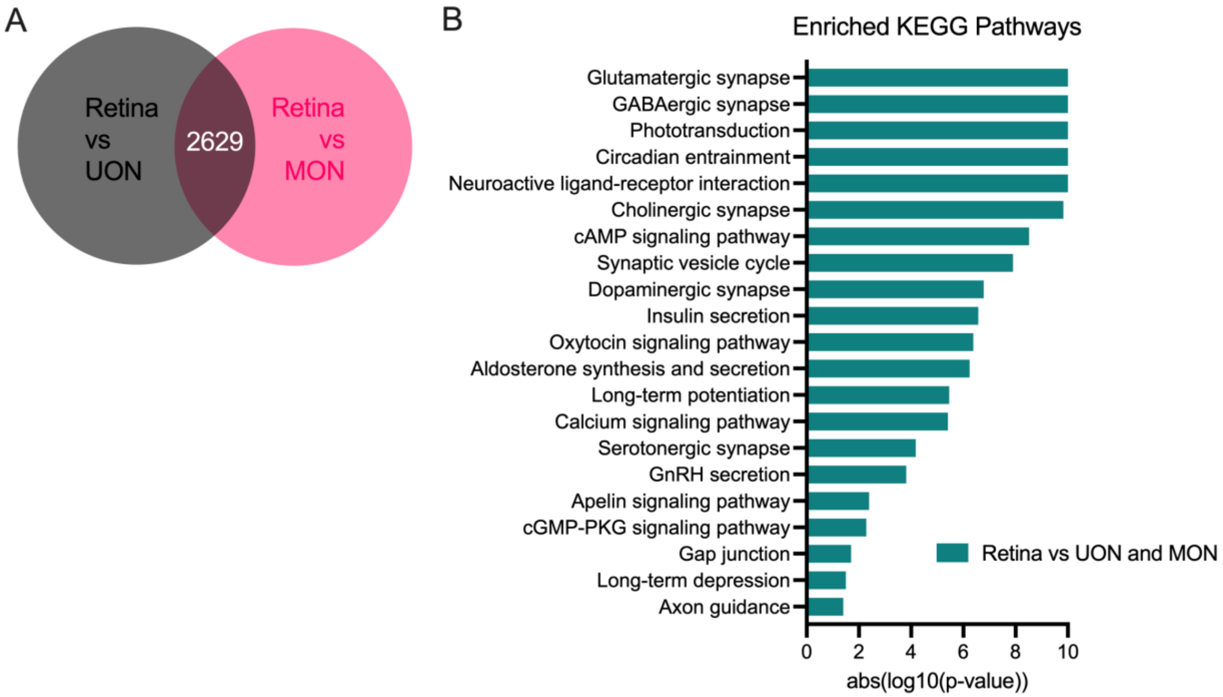


**Figure S2.** Gene signature of the naïve mouse retina. **(A)** Venn diagram showing the number of significantly enriched genes in naïve retinal tissue compared to UON and MON. **(B)** KEGG analysis of enriched retina genes compared to all other tissue regions.

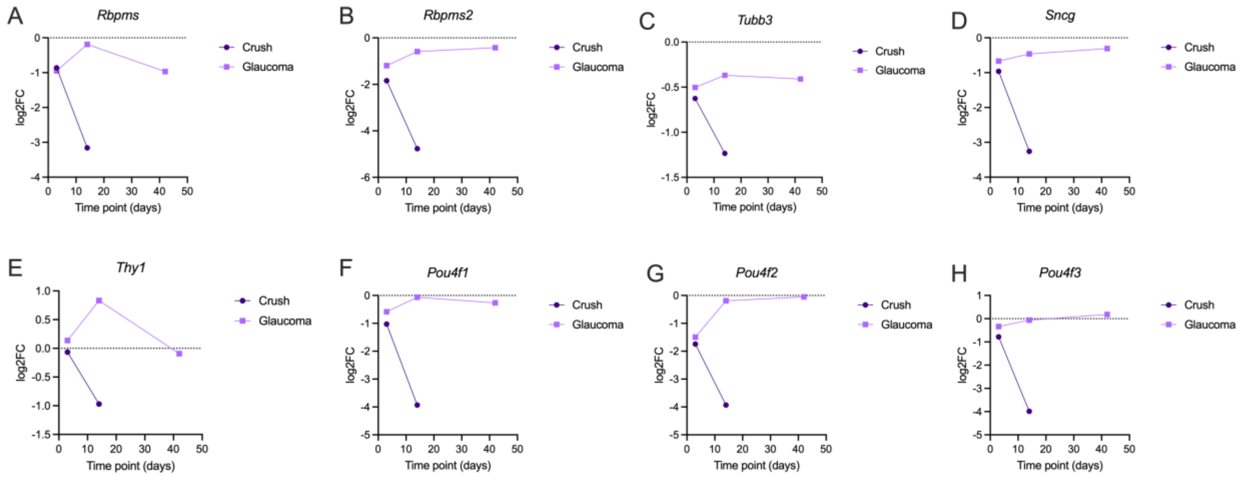


**Figure S3.** Expression of RGC marker genes in retinal tissue following ON injury. **(A-H)** Expression of RGC markers *Rbpms* (A), *Rbpms2* (B), *Tubb3* (C), *Sncg* (D), *Thy1* (E), *Pou4f1* (F), *Pouf4f2* (G), and *Pou4f3* (H) at time points after ON crush or bead-induced glaucoma. Log2FC was calculated from differential expression analysis of each injury time point compared to the naïve 0D control retina samples.

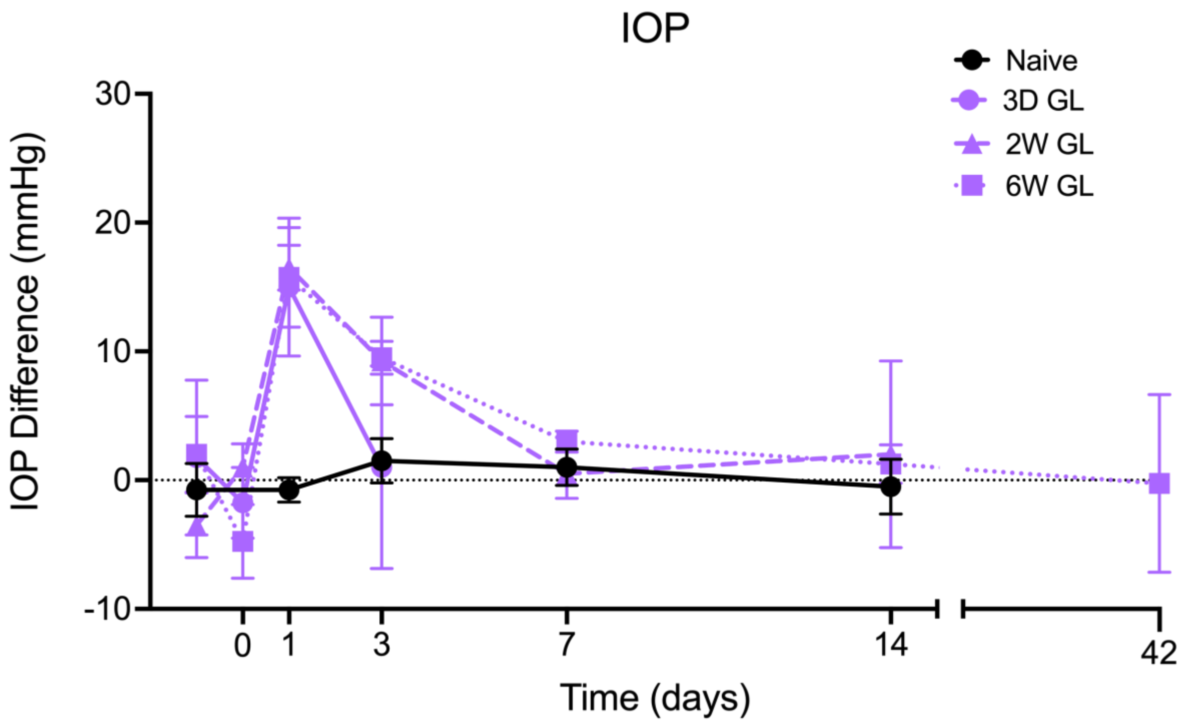


**Figure S4.** IOP measurements over time in the microbead-induced glaucoma model. Difference in IOP (in mmHg) between microbead-injected and contralateral control eyes over time in mice used in the glaucoma model RNA-seq study. Each line represents IOP measurements from the group of animals used for each collection time point, and error bars indicate standard deviation.

**Table S1.** IOP difference measurements (mmHg) over time in microbead-induced glaucoma animals.

| | | Mean IOP difference (in mmHg) (left) and Stdev (right) | | | | | | | |
|---|---|---|---|---|---|---|---|---|---|
| | | Naïve (0D) | | 3D GL | | 2W GL | | 6W GL | |
| IOP Time point (pre/post-bead and days following) | Pre-bead | -0.75 | 2.06 | 1.75 | 6.02 | -3.5 | 2.52 | 2 | 2.94 |
| | Post-bead | | | -1.75 | 2.75 | 1 | 1.83 | -4.75 | 2.87 |
| | 1 | -0.75 | 0.96 | 15 | 5.35 | 16.5 | 1.73 | 15.75 | 3.86 |
| | 3 | 1.5 | 1.73 | 1 | 7.87 | 9.25 | 3.4 | 9.5 | 1.29 |
| | 7 | 1 | 1.41 | | | 0.5 | 1.91 | 3 | 0.82 |
| | 14 | -0.5 | 2.12 | | | 2 | 7.26 | 1.25 | 1.5 |
| | 30 | | | | | | | -0.25 | 6.9 |

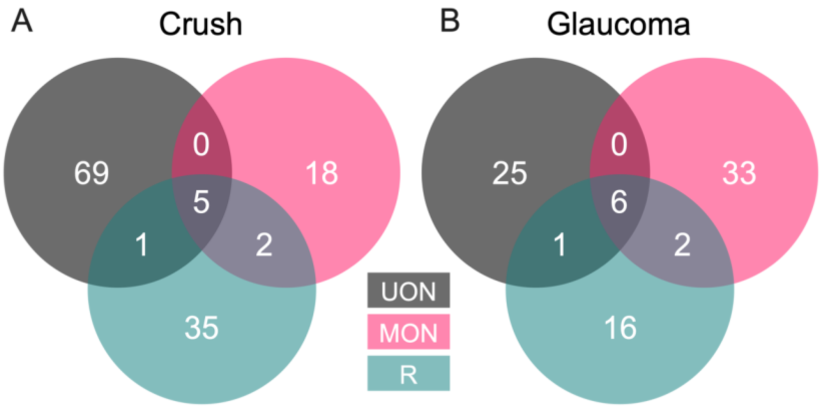


**Figure S5.** Sex-specific gene expression differences in disease models. **(A-B)** Venn diagrams showing the numbers of DEGs in differential expression analysis comparing male and female replicates for each tissue region in crush (A) and glaucoma (B) samples. Male and female replicates in all time points for each model were analyzed collectively.

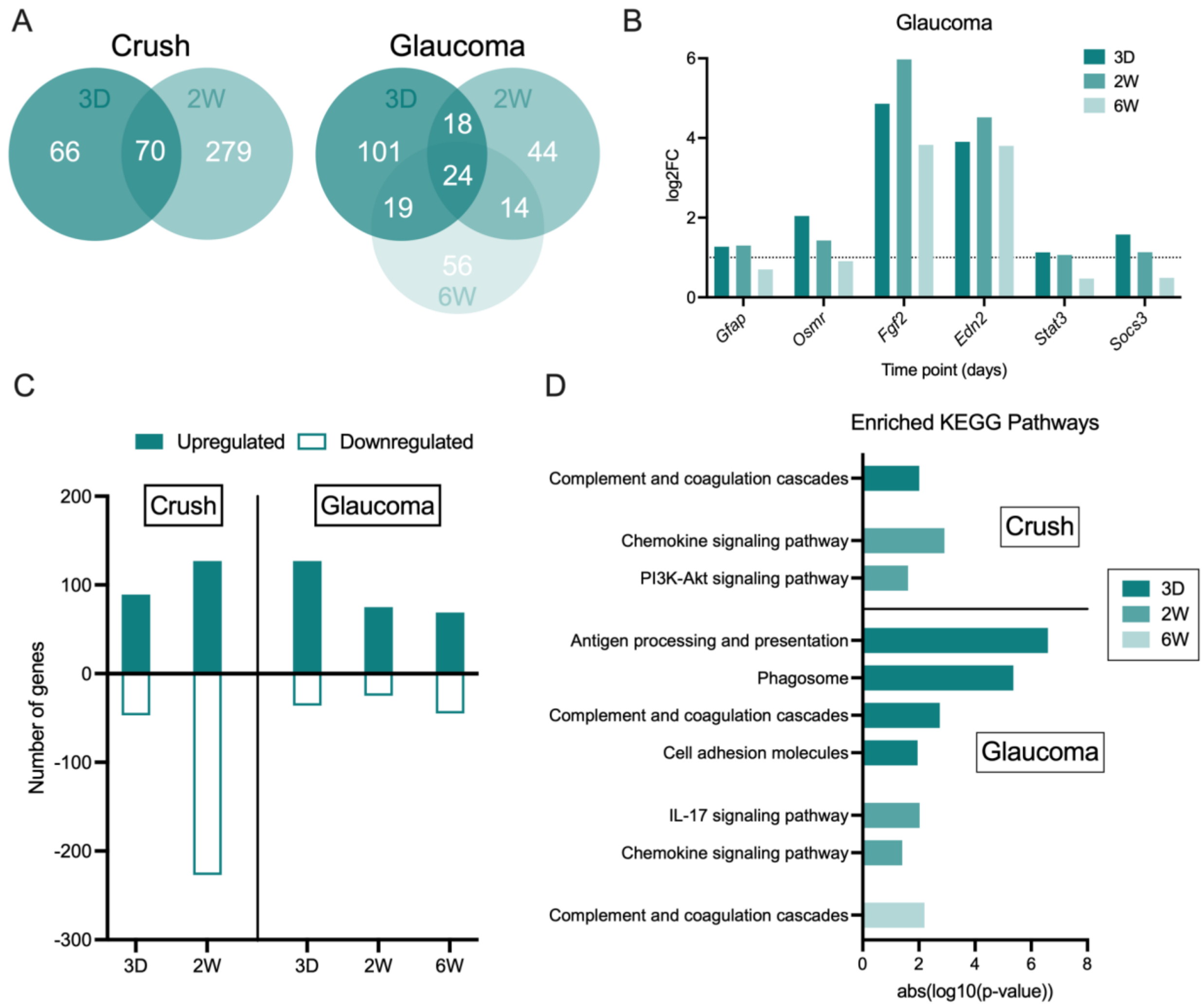


**Figure S6.** Differential responses to crush and glaucoma in the retina. **(A)** Venn diagrams showing relationships of retinal tissue gene responses to ON crush (left) and bead-induced glaucoma (right). **(B)** Upregulation of stress response genes in the retina at early time points of the glaucoma model. Dotted line indicates significance threshold of log2FC > 1. **(C)** Number of up/down genes in retinal tissue at each crush and glaucoma time point. **(D)** KEGG pathway analysis of UON and MON DEGs at different time points following ON crush and IOP elevation.

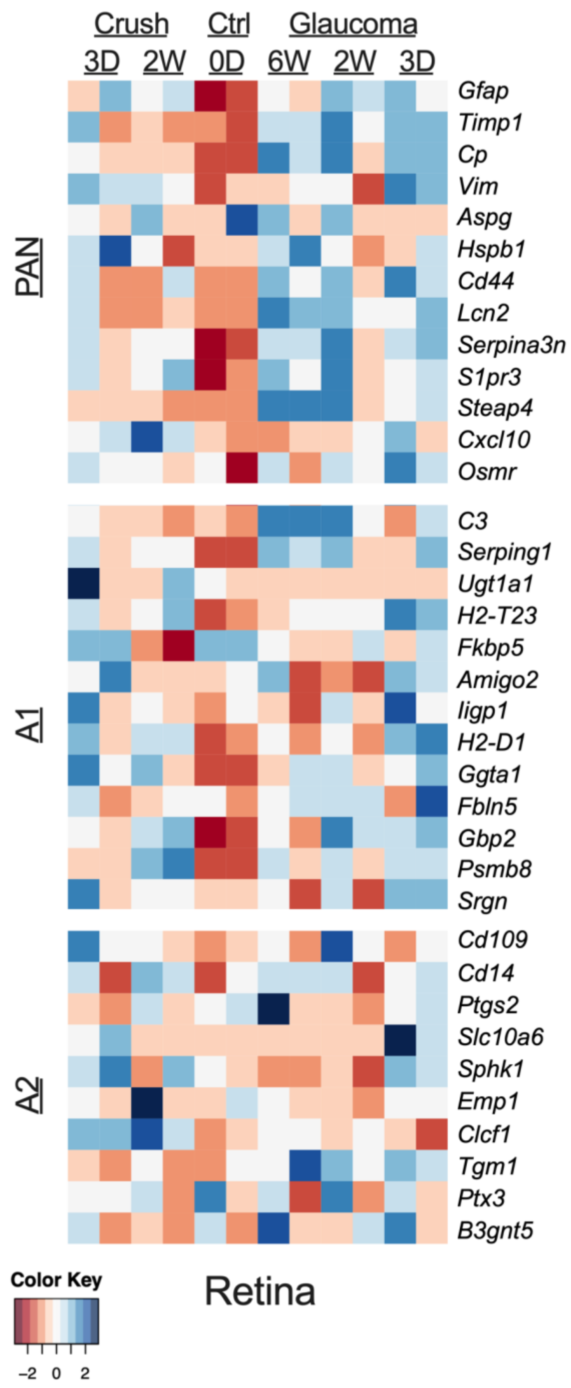


**Figure S7.** A1/A2-specific gene signatures in the retina. Heatmap showing PAN-reactive, A1-specific, and A2-specific astrocyte markers in naïve and injured retinal tissue.

**Table S2:** Animals used in this study.

| Strain | Treatment(s) | Time Point(s) | N Animals (+ Sex) | Eye (OS/OD) |
|---|---|---|---|---|
| C57BL/6 (B6) | Naïve | 0D | 4 (2M, 2F) | OS |
| | ON Crush | 3D | 4 (2M, 2F) | OS |
| | | 2W | 4 (2M, 2F) | OS |
| | Glaucoma | 3D | 4 (2M, 2F) | OS |
| | | 2W | 4 (2M, 2F) | OS |
| | | 6W | 4 (2M, 2F) | OS |
| Wild-type (non-fluorescent) FVB/N-Tg(GFAP-GFP)14Mes littermate mice (FVB or WT-GFP) | Naïve | 0D | 3F | OS and OD |

**Table S3.** qPCR primers used in this study.

| Gene | Forward Sequence (5' - 3') | Reverse Sequence (5' - 3') |
|---|---|---|
| *Actb* | ACCTTCTACAATGAGCTGCG | CTGGATGGCTACGTACATGG |
| *Gapdh* | CCAATGTGTCCGTCGTGGATC | GCTTCACCACCTTCTTGATGTC |
| *Rpl19* | TCACAGCCTGTACCTGAA | TCGTGCTTCCTTGGTCTTAG |
| *Gfap* | CAGAGGAGTGGTATCGGTCTAA | GATAGTCGTTAGCTTCGTGCTT |
| *Ptx3* | GACTTCATCCCACCGAGGAC | CATGCCCTCCCGCATCT |
| *Sphk1* | GGAGACTGCCATCCAGAAAC | GGTCTTCATTAGTCACCTGCTC |
| *Stat3* | TGCTGCCCCGTACCTGAAGA | GGACATCGGCAGGTCAATGGTATTG |
| *C3* | TGGAGAAAGCAGTGATGGTAAG | GTCCACAGTGAAGATCCGATATAA |
| *Serping1* | ACAGCCCCCTCTGAATTCTT | GGATGCTCTCCAAGTTGCTC |
| *C1qa* | TGGAGCATCCAGTTTGATCG | TGTCCATACTAGGGTCATGGT |
| *Il1a* | GCTTGAGTCGGCAAAGAAATC | GAGAGATGGTCAATGGCAGAA |
| *Ptk2 (FAK)* | CGATGAGGAAGACACATACACC | TCCAAACTGACCTTCTCCAATAC |
| *Rock1* | CAGACCTCACAGCTTGCTAAT | GCTCATCTCTGTGTGACTCTTC |
| *Rock2* | GAGATGAGTGCAGCAGCTATTA | TCATGATCTCCGCCAACTTATT |
| *Piezo2* | GGGCACCTGATTGGACTTTA | GTCTGTCTGGATAACGGACTTG |
| *Sox2* | AACGGCAGCTACAGCATGATGC | CGAGCTGGTCATGGAGTTGTAC |

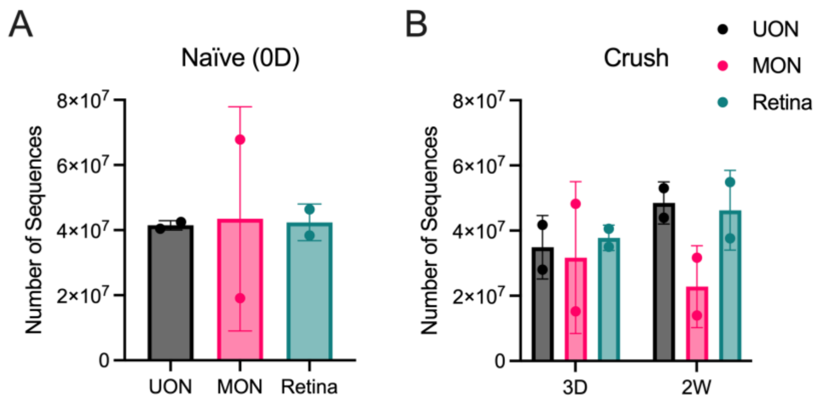


**Figure S8.** Total number of sequences. **(A-C)** Number of sequences for each naïve (A), ON crush (B), and glaucoma model (C) RNA-seq samples included in the study. Dots represent number of sequences for a single, pooled, replicate and error bars indicate standard deviation. 3D, three days; 2W, two weeks; 6W, six weeks.
